## Supplementary figures and images for "A single-cell atlas of the miracidium larva of the human blood fluke *Schistosoma mansoni*: cell types, developmental pathways and tissue architecture"

### Sup Fig 2.tif

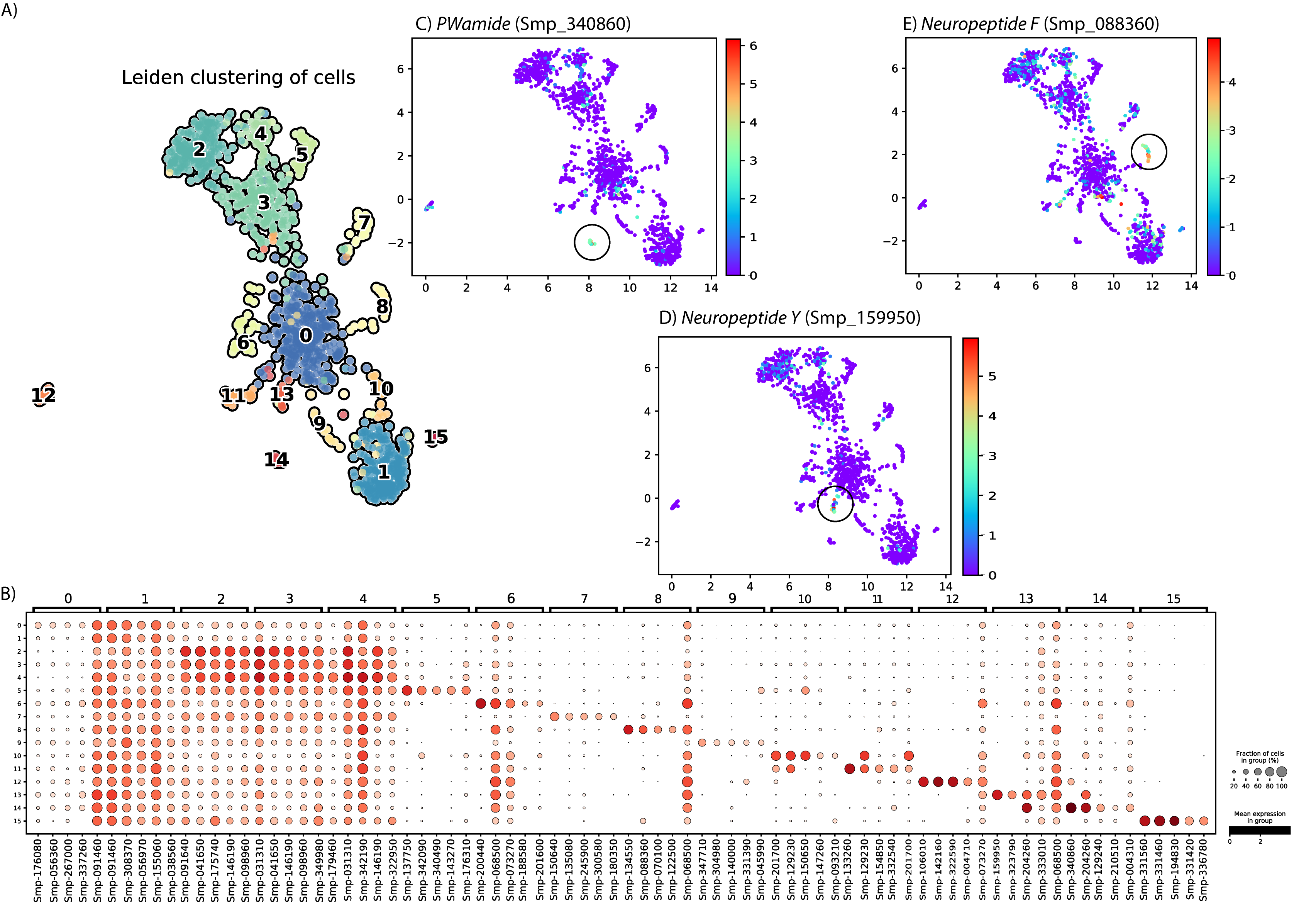

### Sup Fig 4.tif

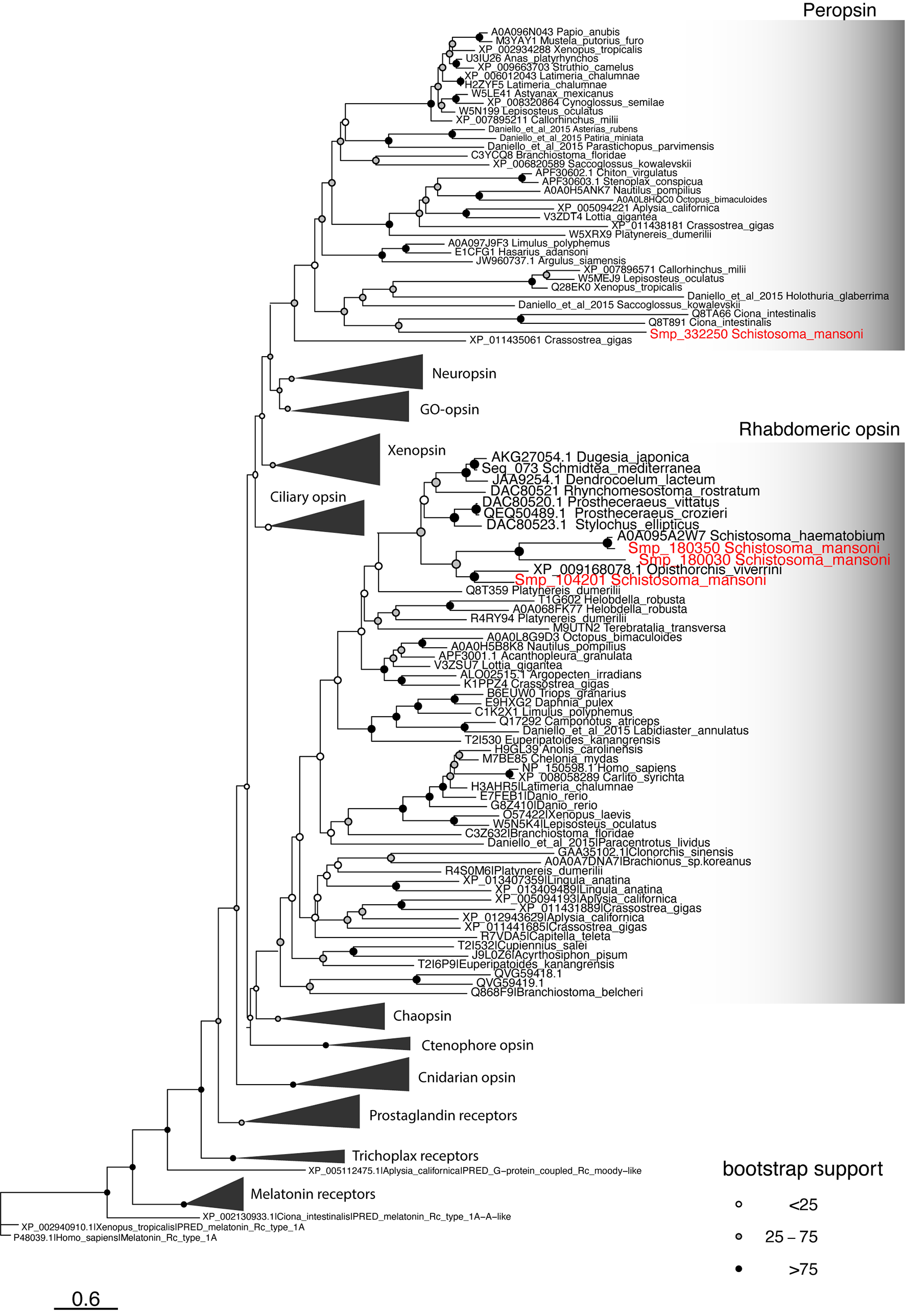

### Sup Fig 5.tif

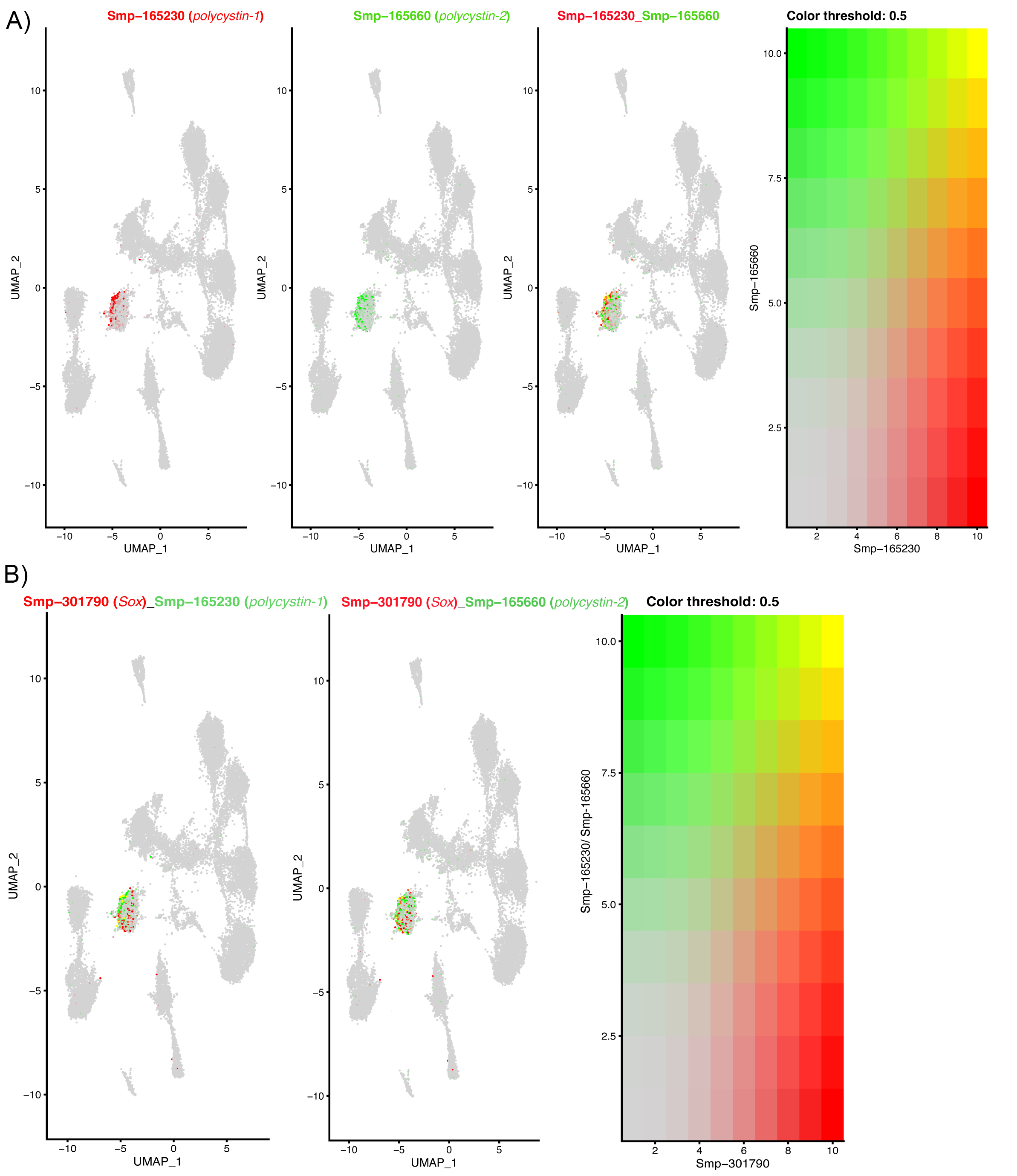

### Sup Fig 6.tif

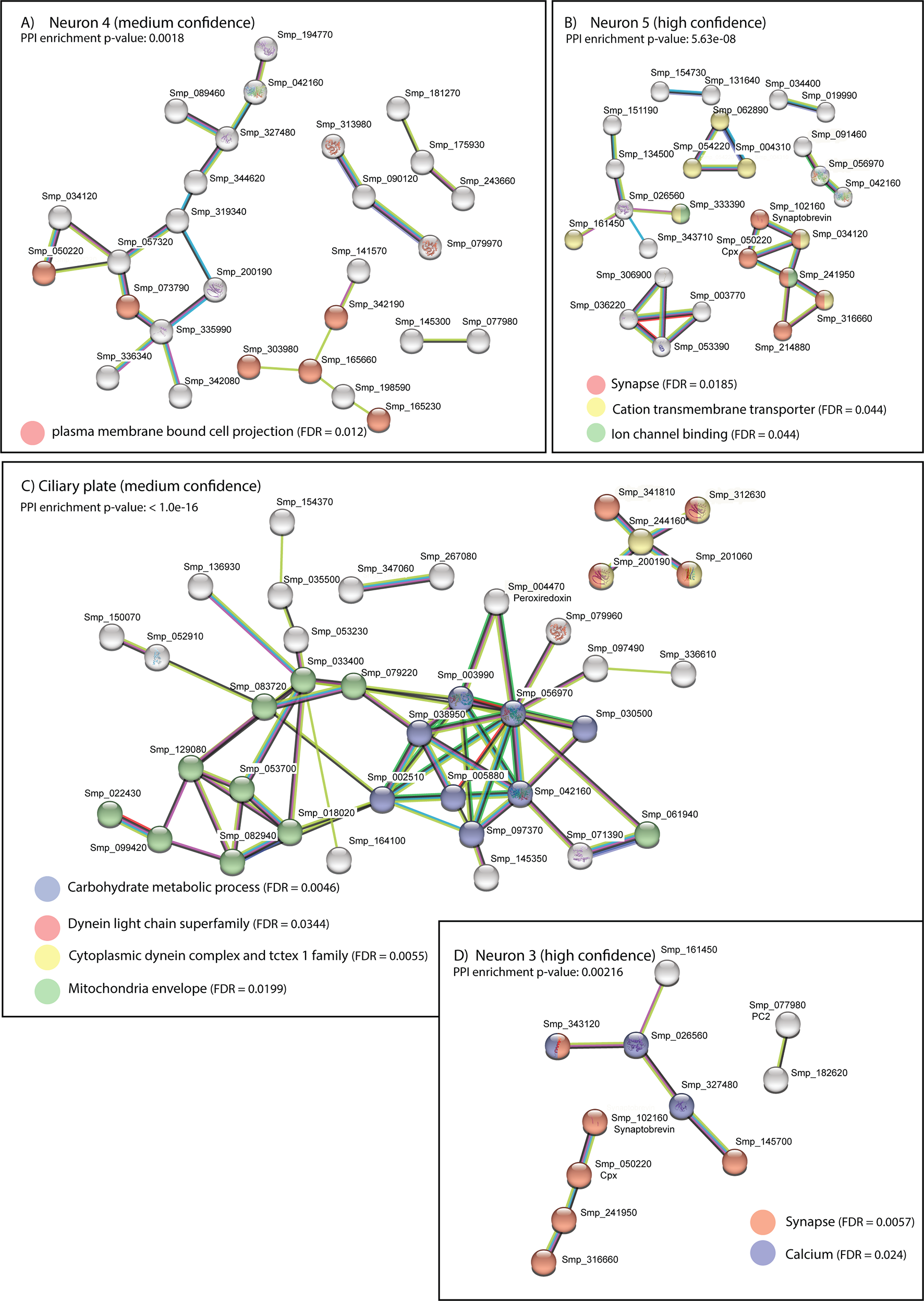

### Sup Fig 7.tif

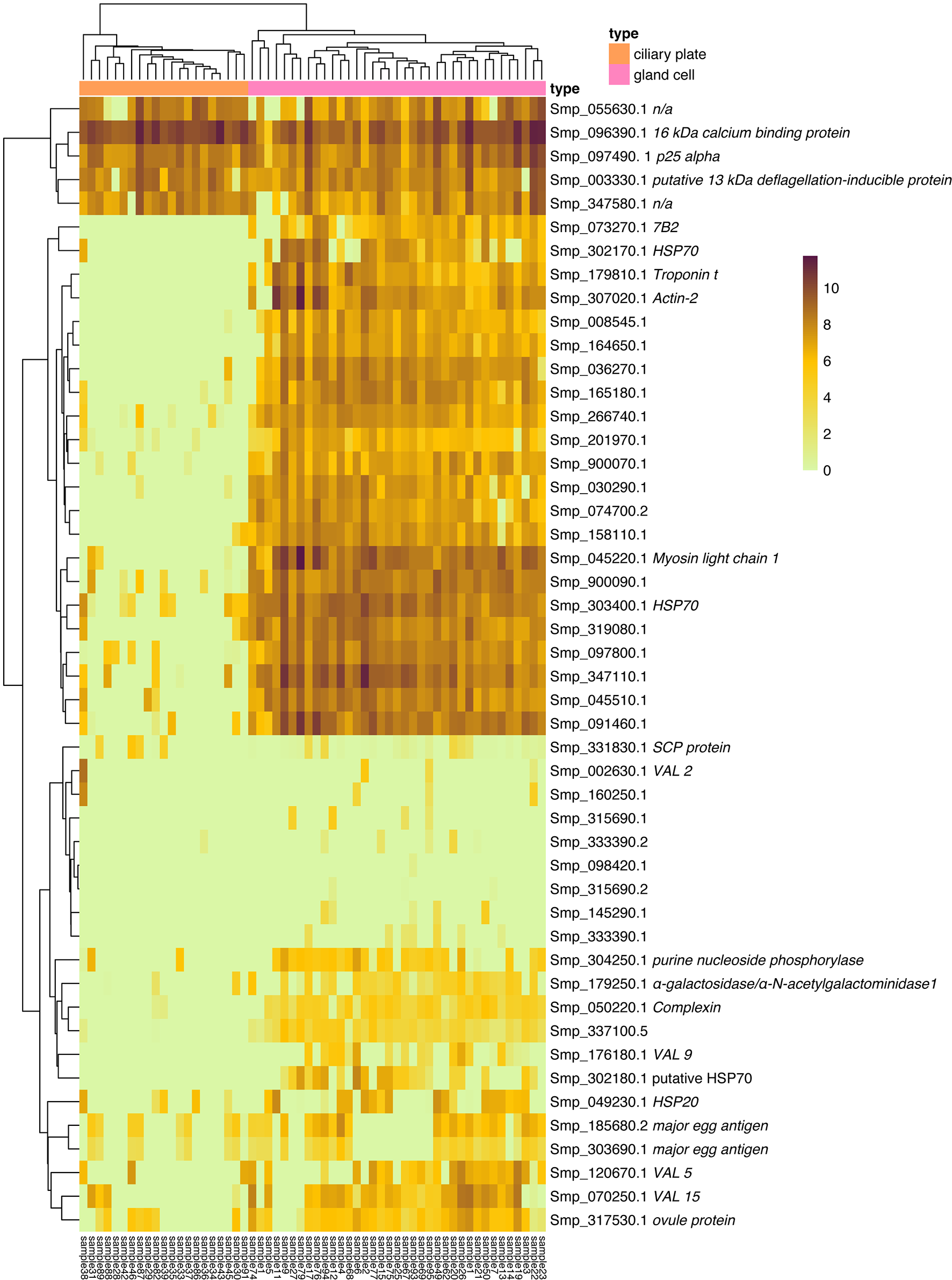

### Sup Fig 8.tif

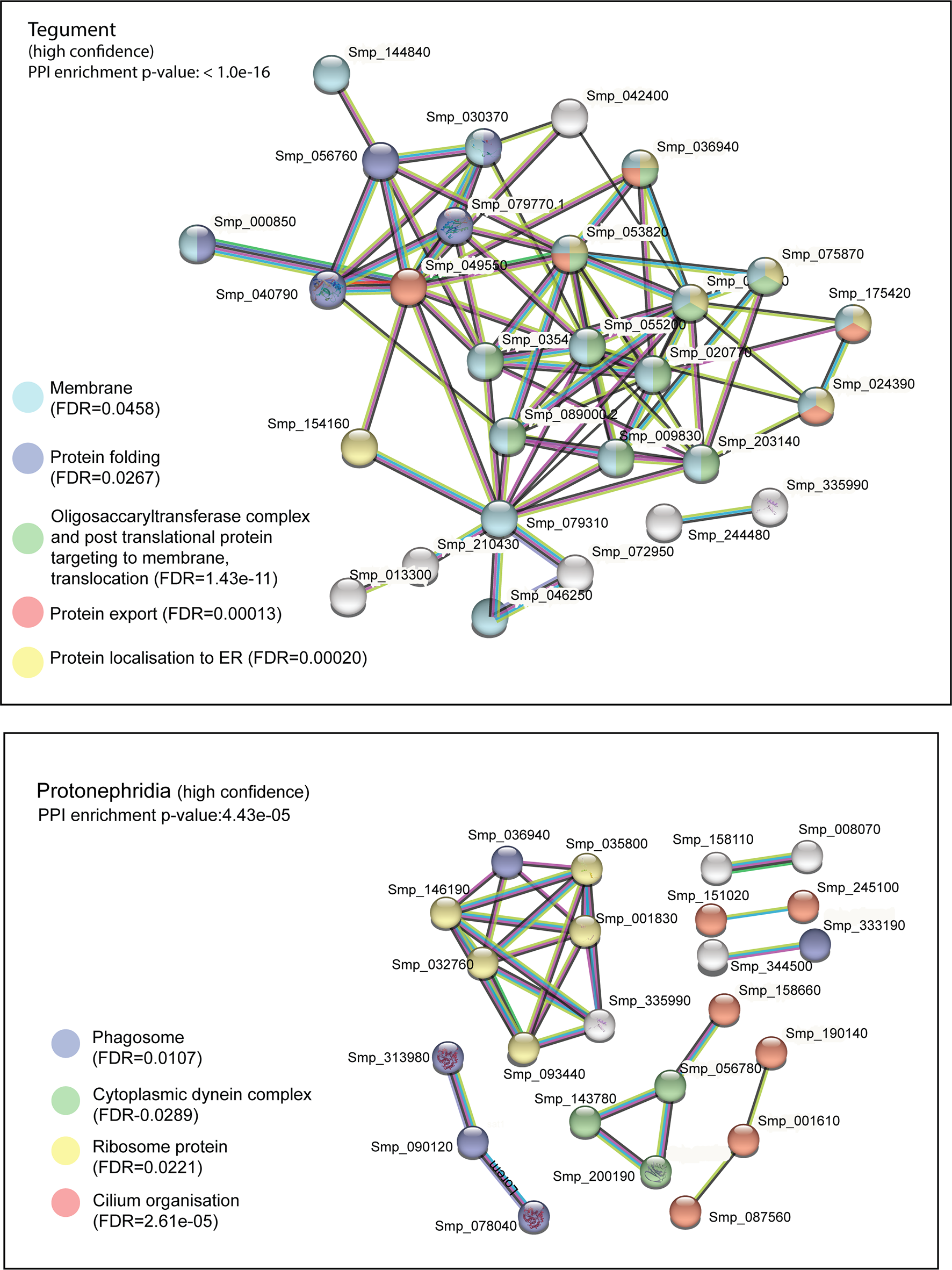

### Sup Fig 9.tif

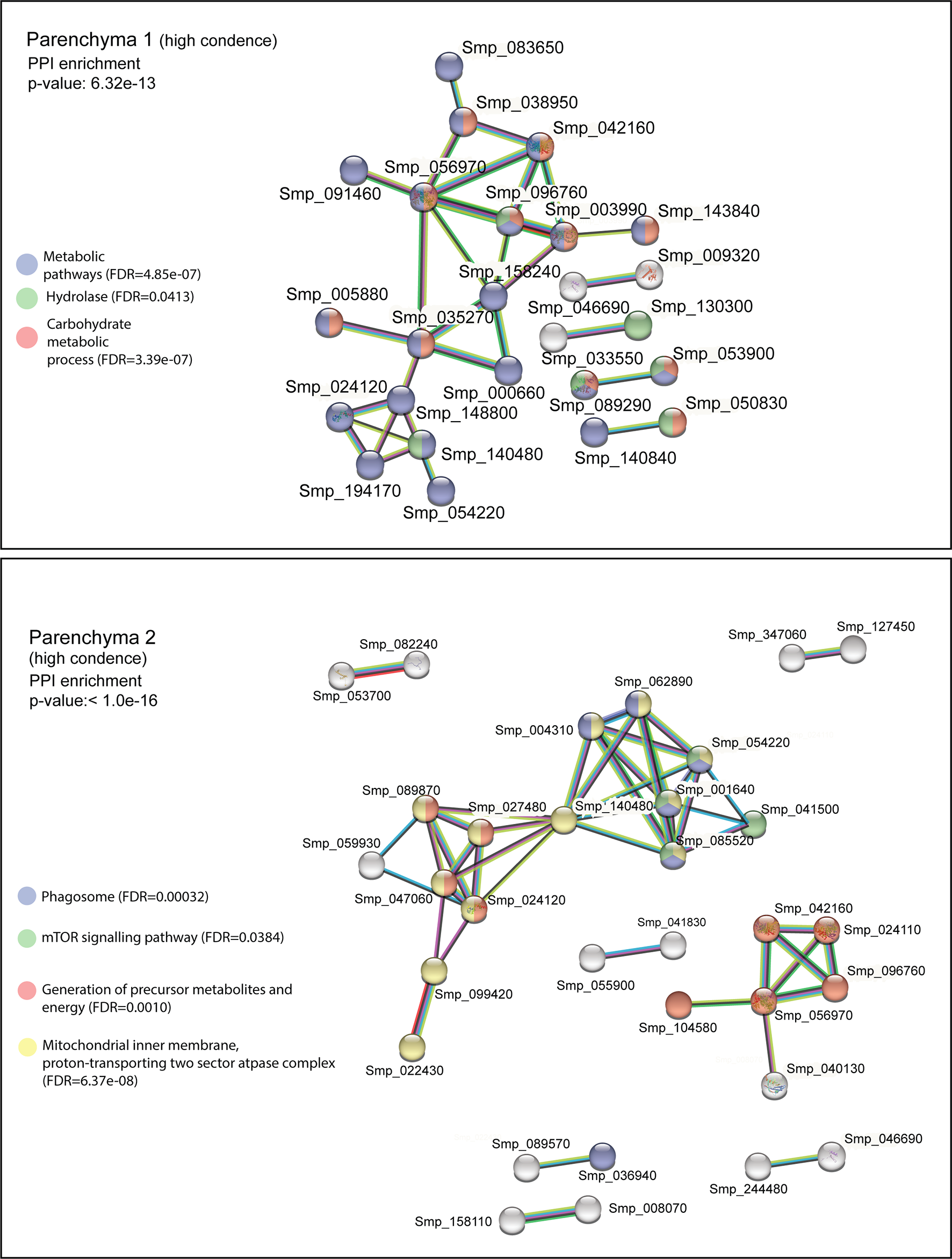

### Sup Fig 10.tif

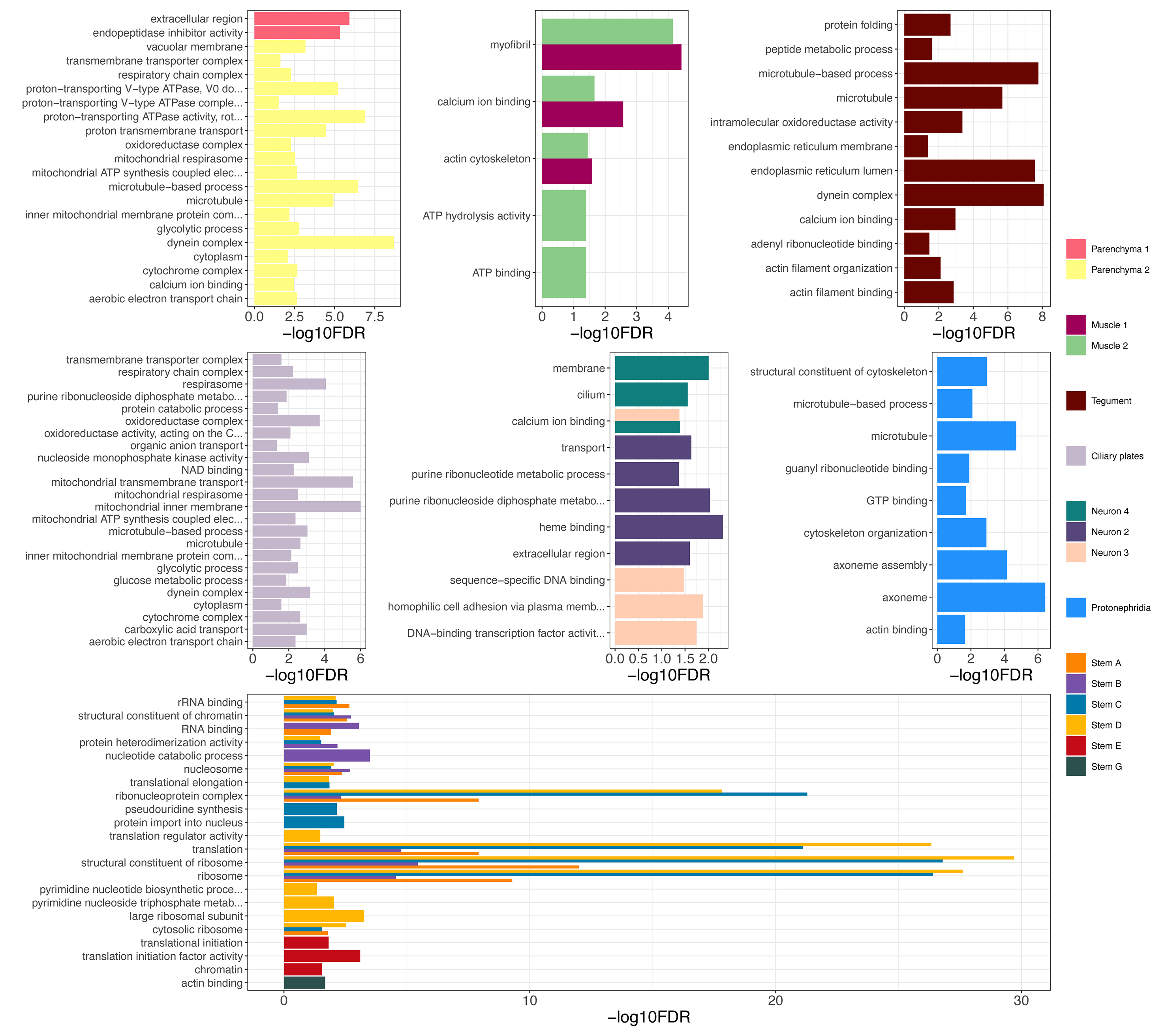

### Sup Fig 12.tif

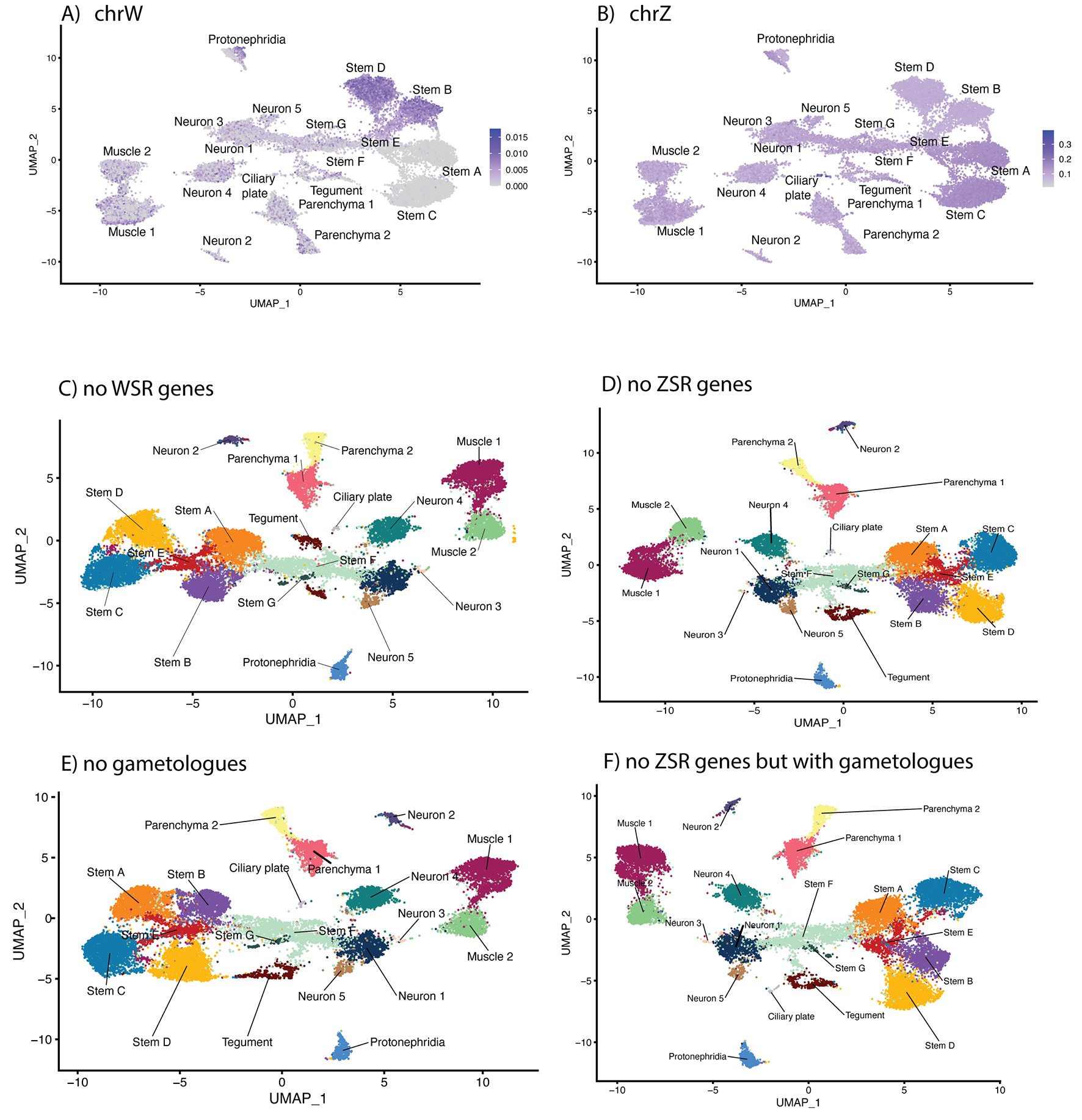

### Sup Fig 14 ZW dosage.tif

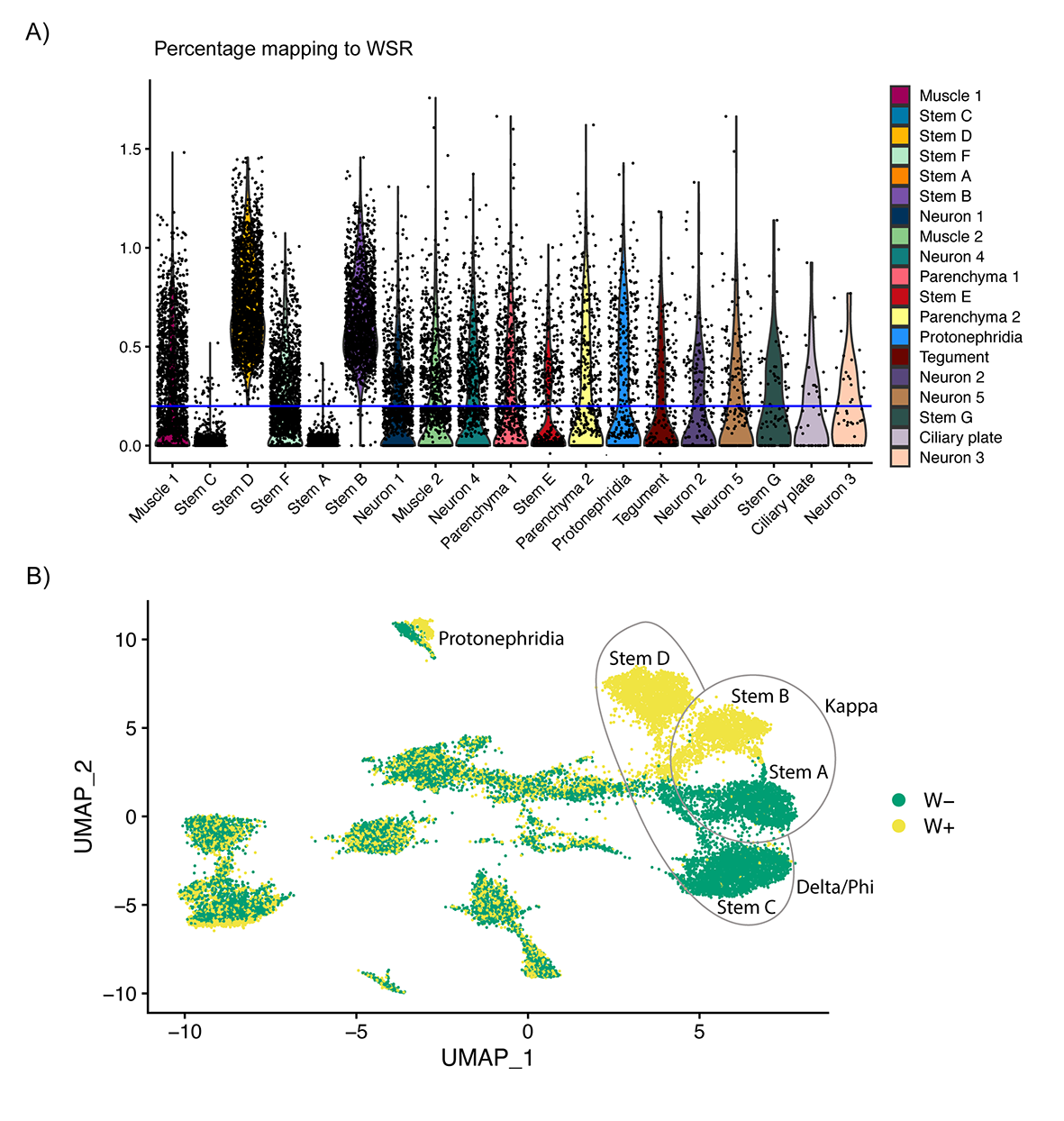

### Sup Fig 15.tif

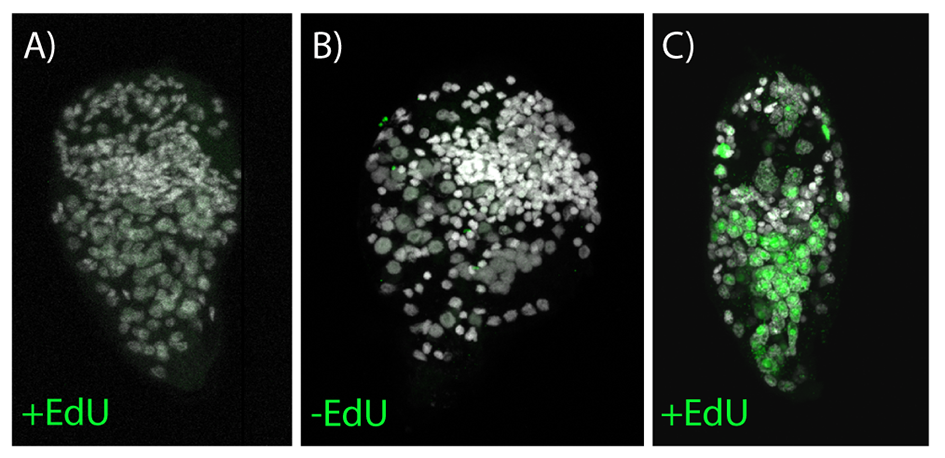

### Sup Fig 16.tif

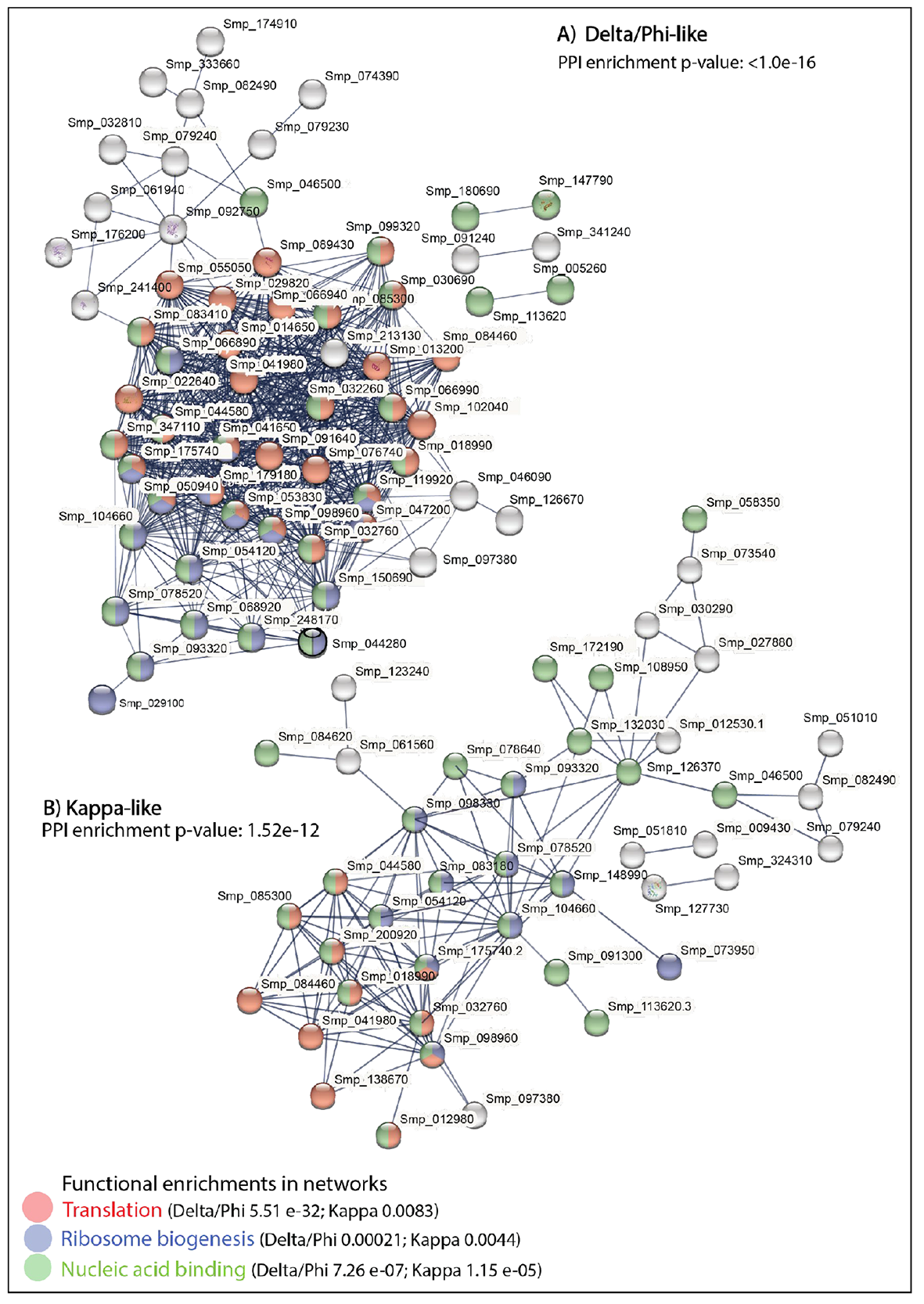

### Sup Fig 17.tif

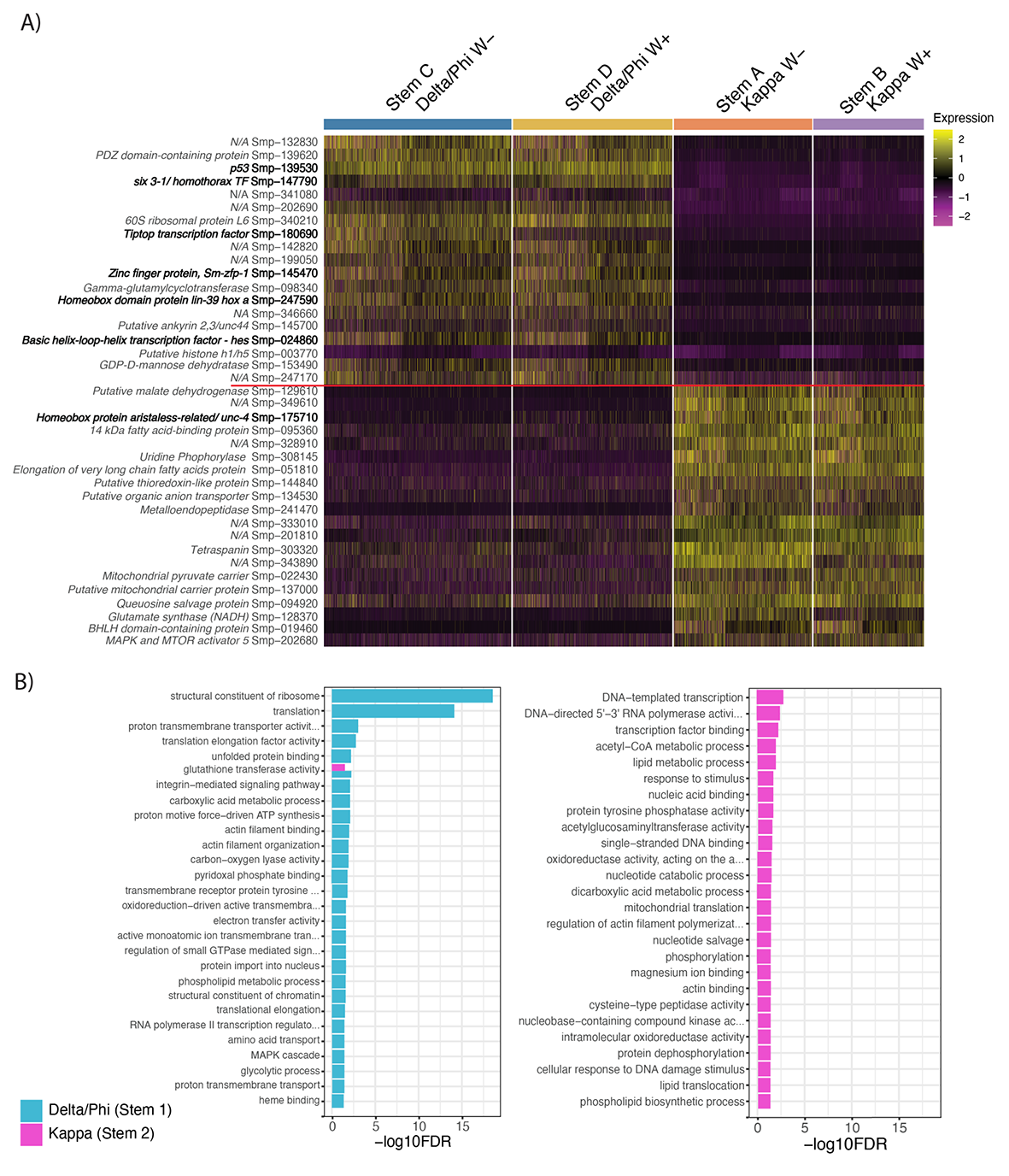

### Sup Fig 18.tif

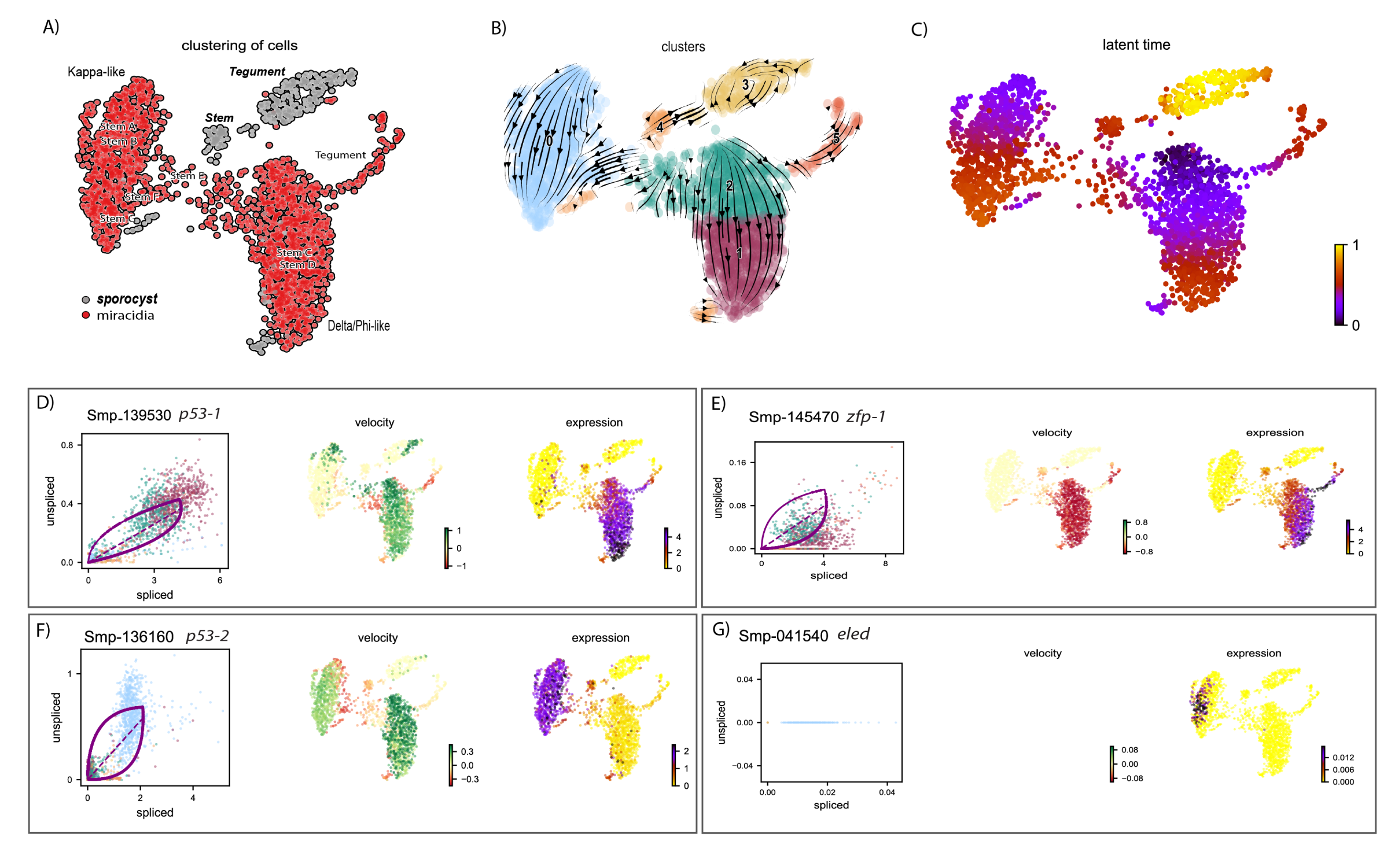

### Sup Fig 19.tif

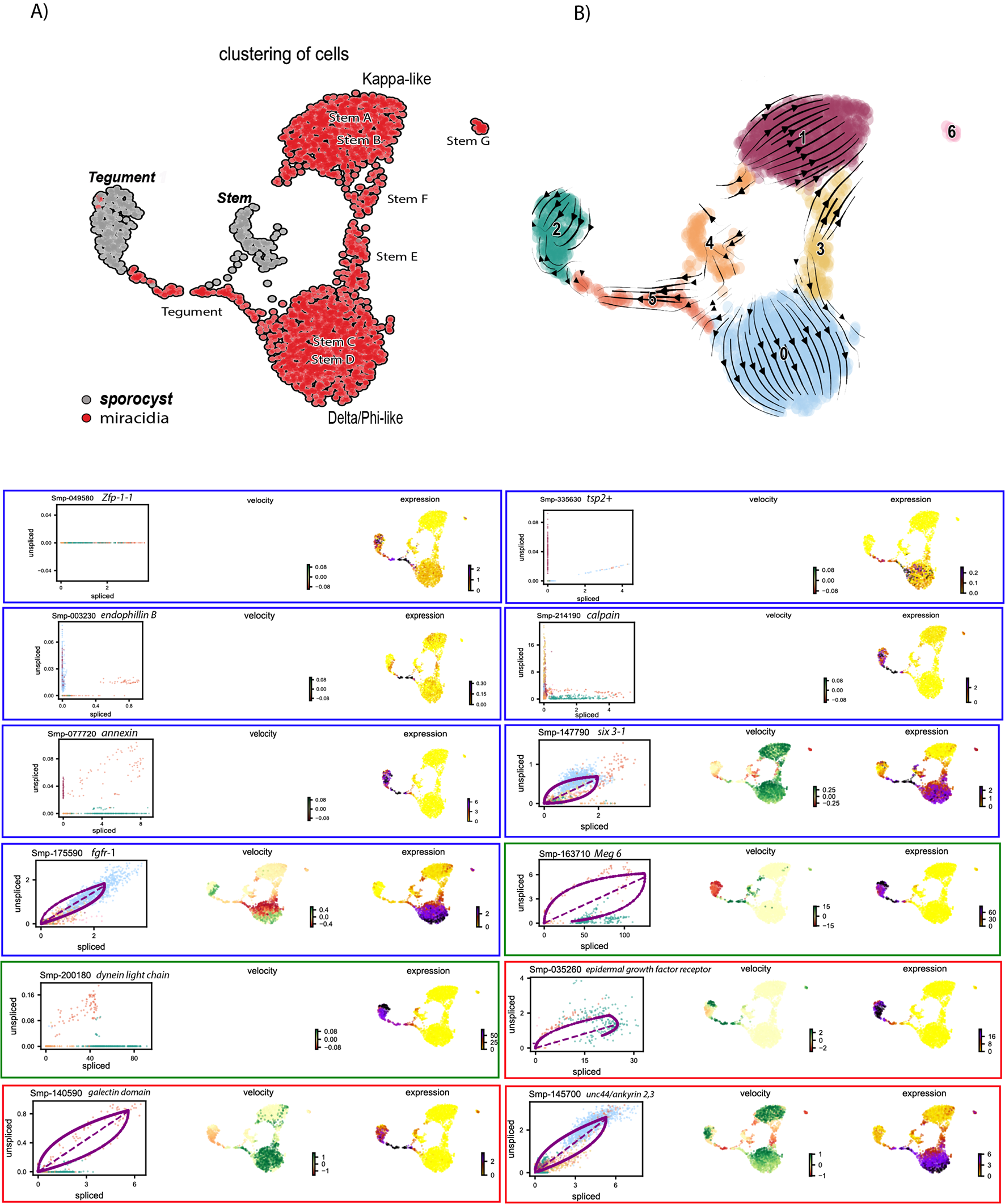

### Sup Fig 20.tif

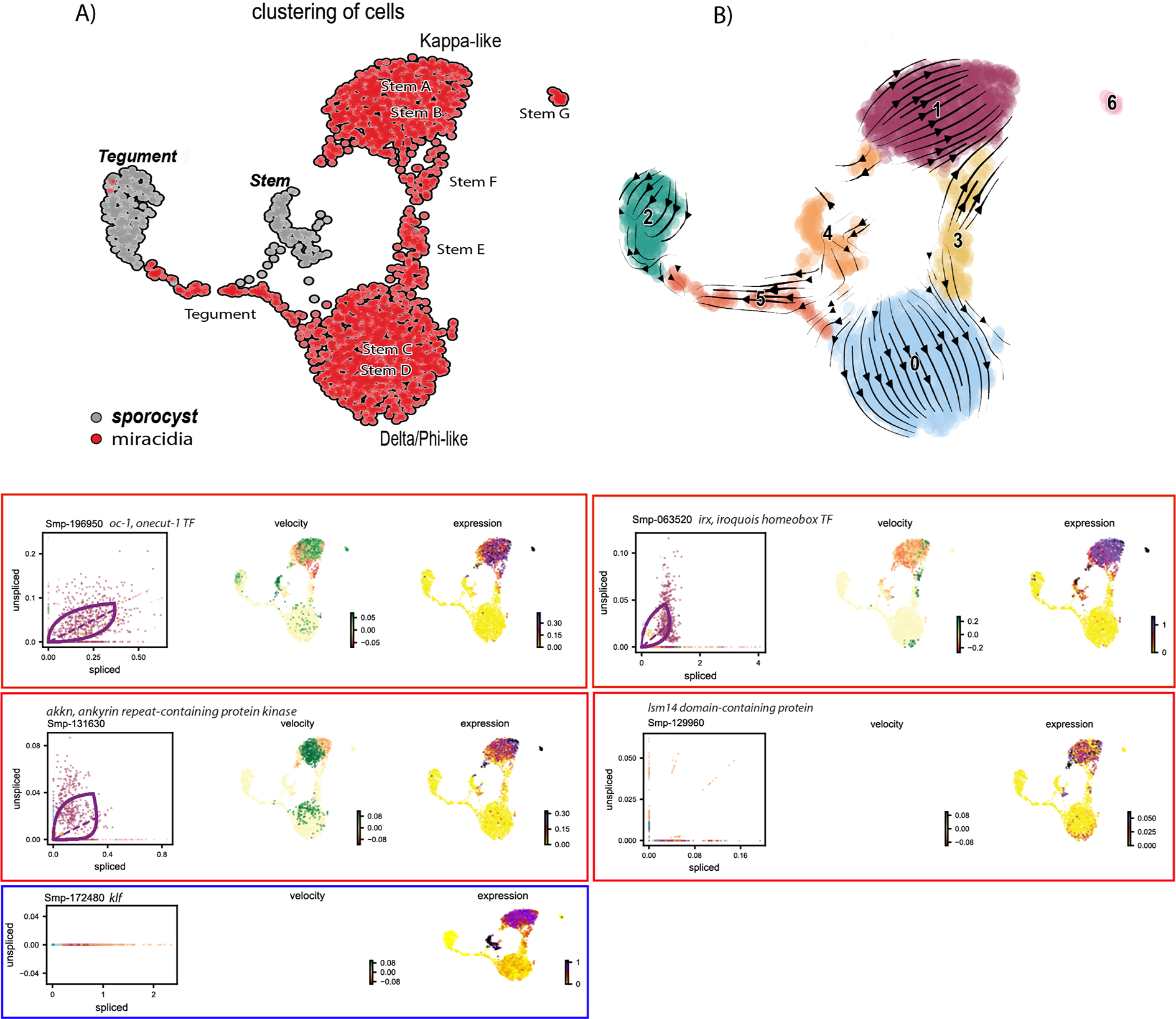

### Sup Fig 21.tif

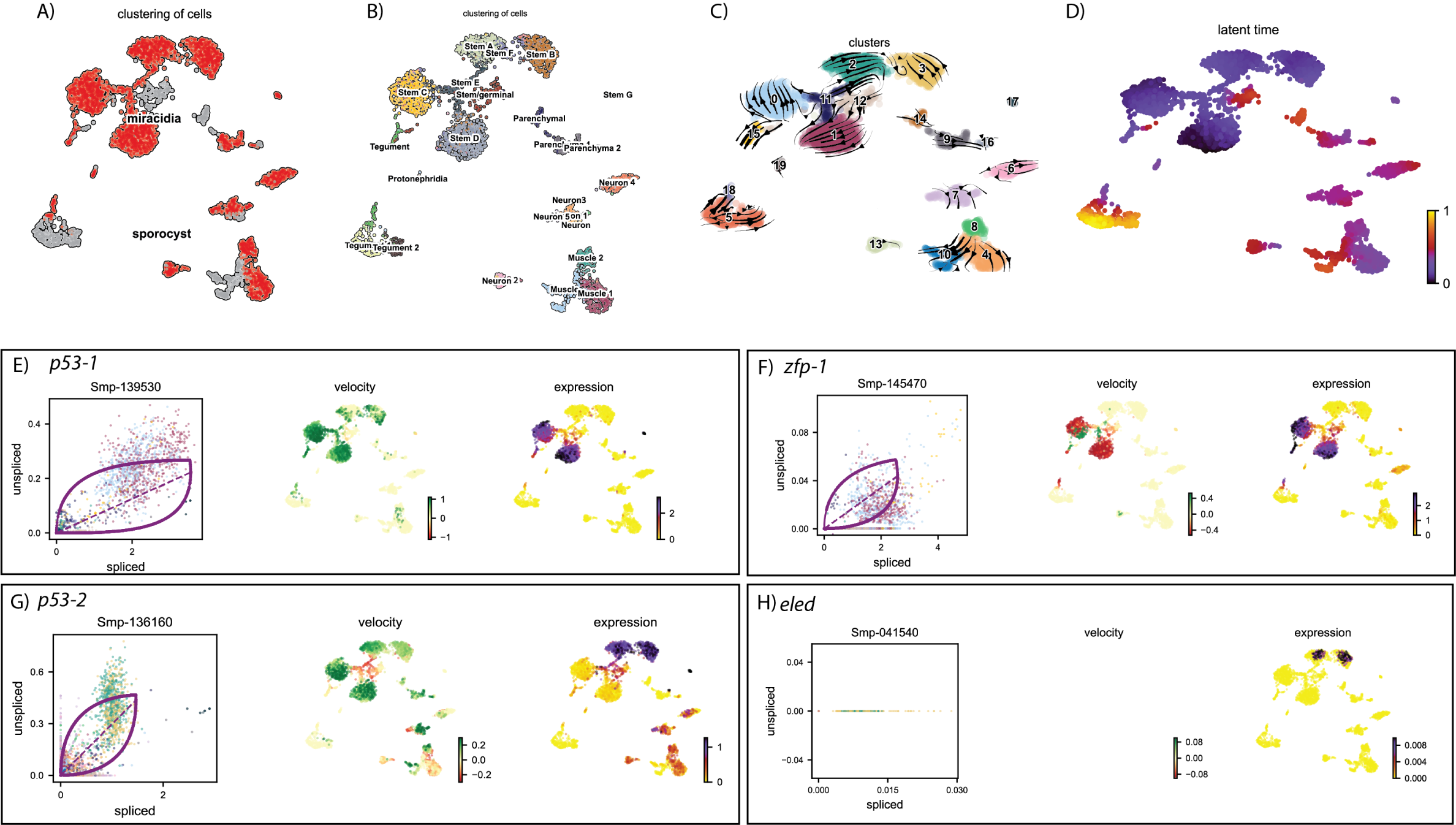

### Sup Fig 22.tif

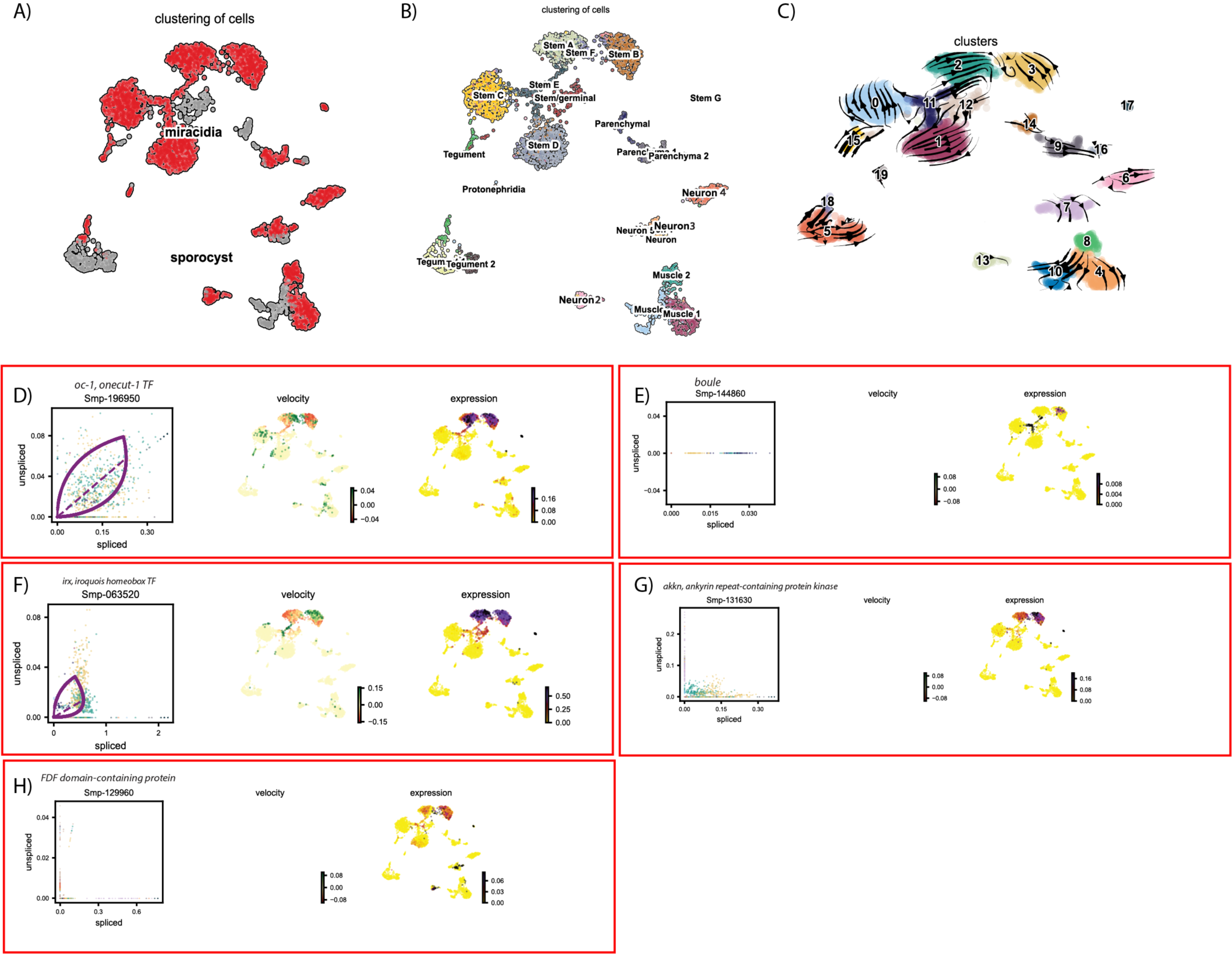

### Sup Fig 23.tif

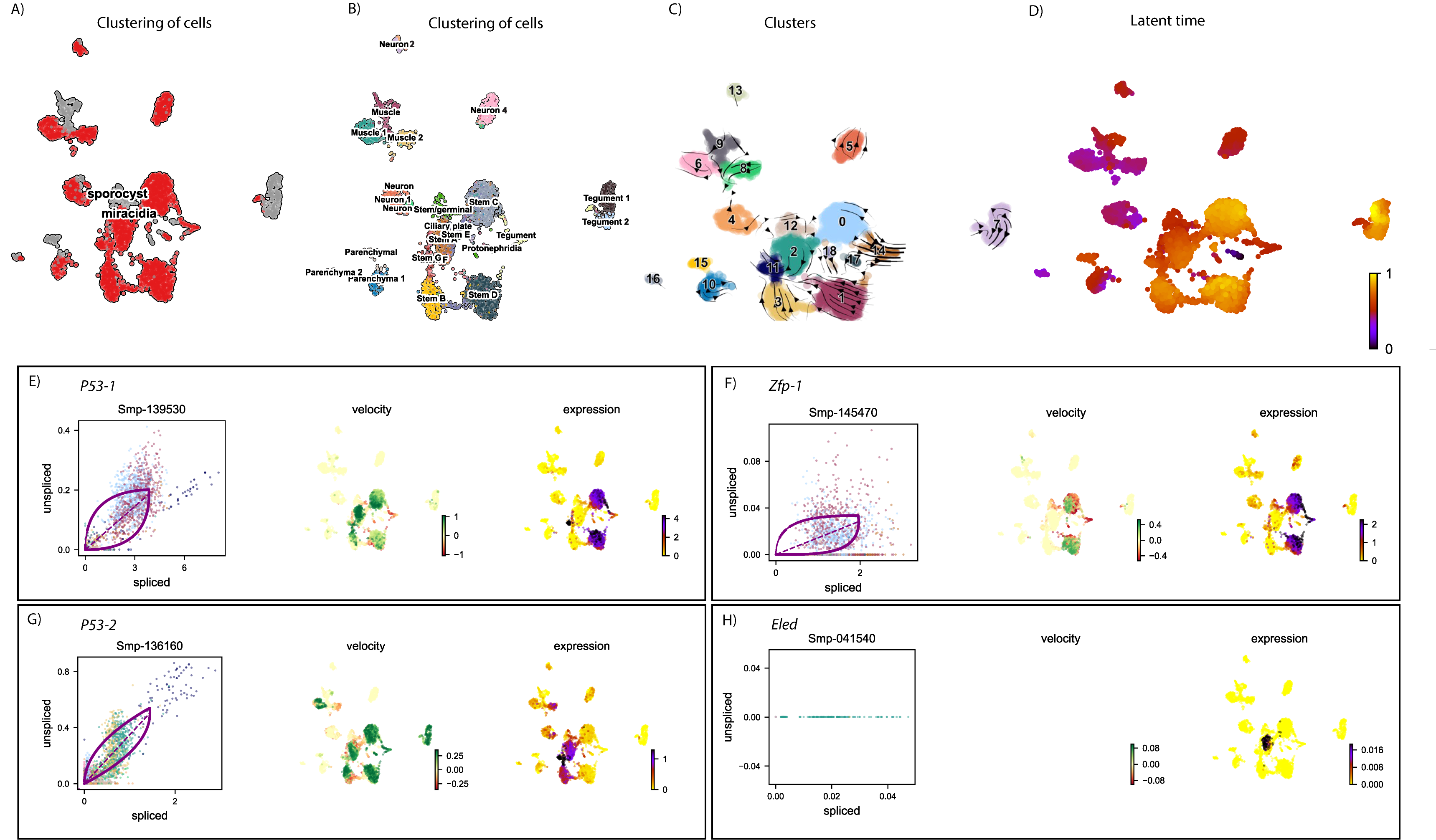

### Supp Fig 1.tif

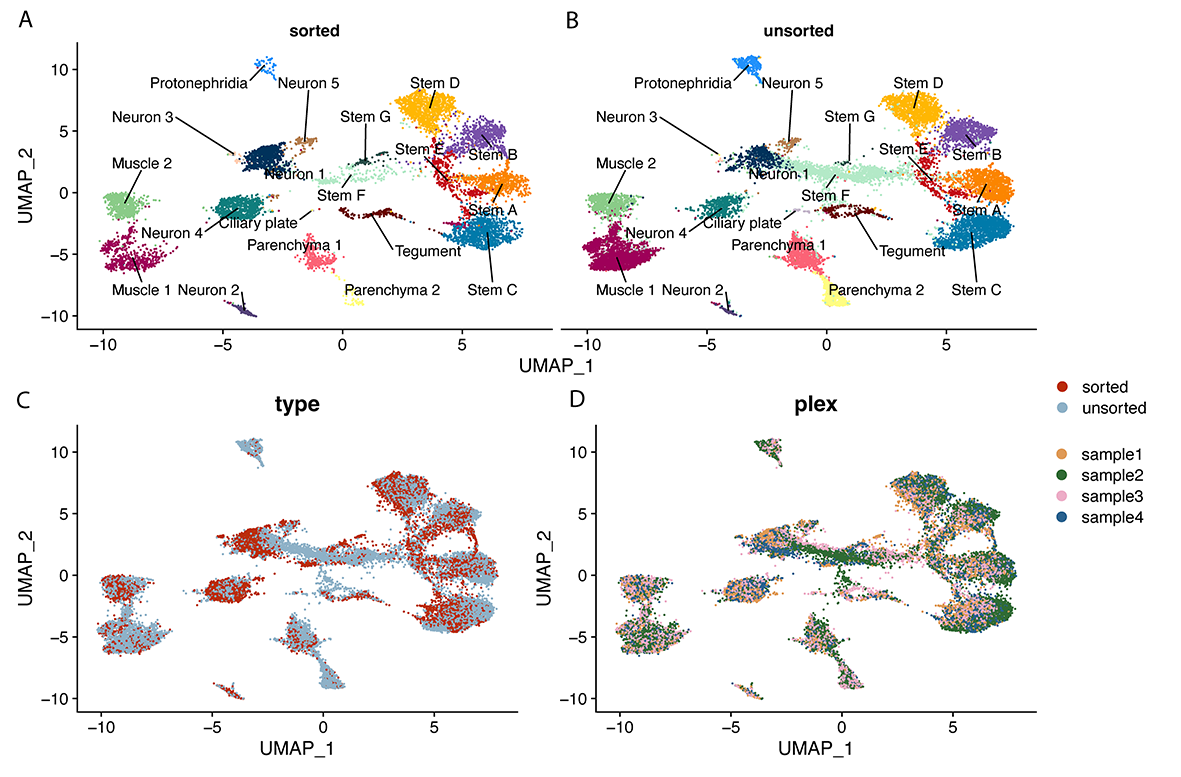

### Supp Fig 3.tif

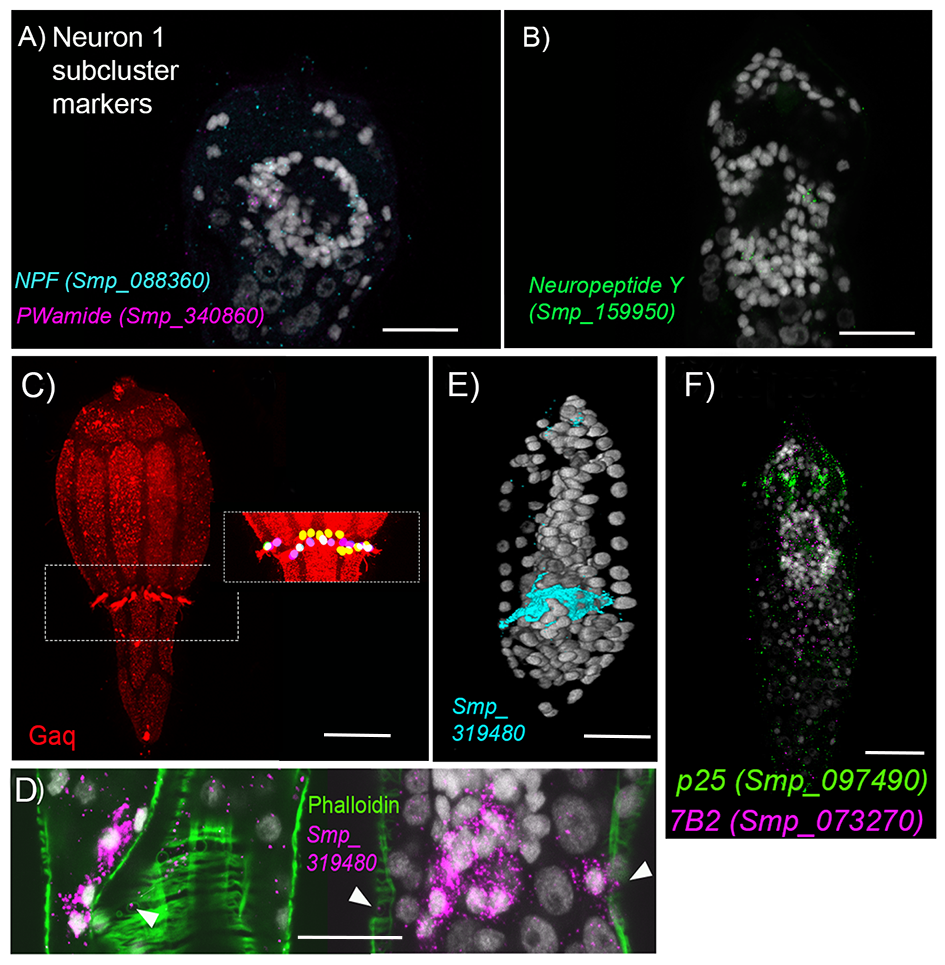

### Supp Fig 11.tif

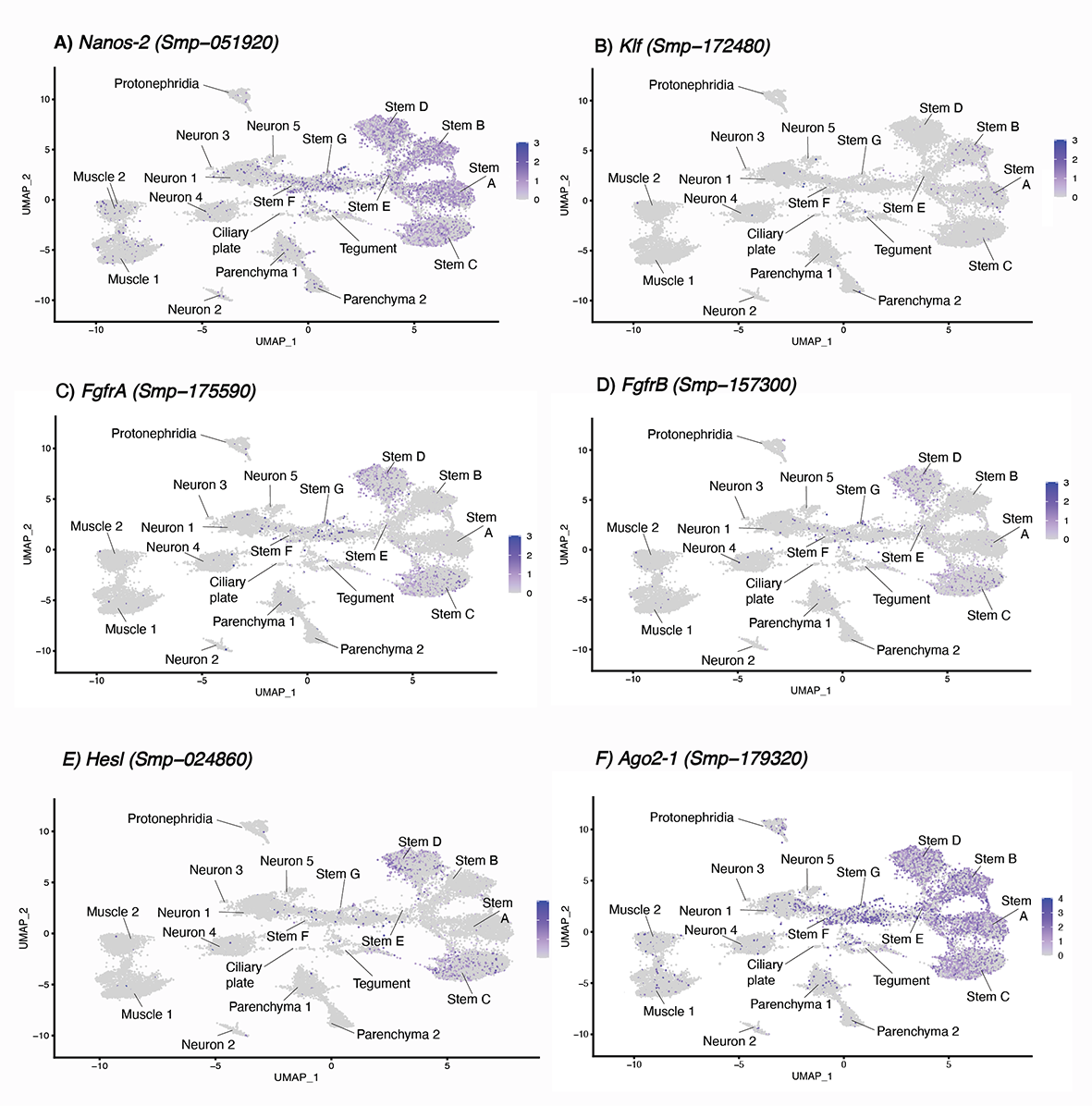

### Supp Fig 13.tif

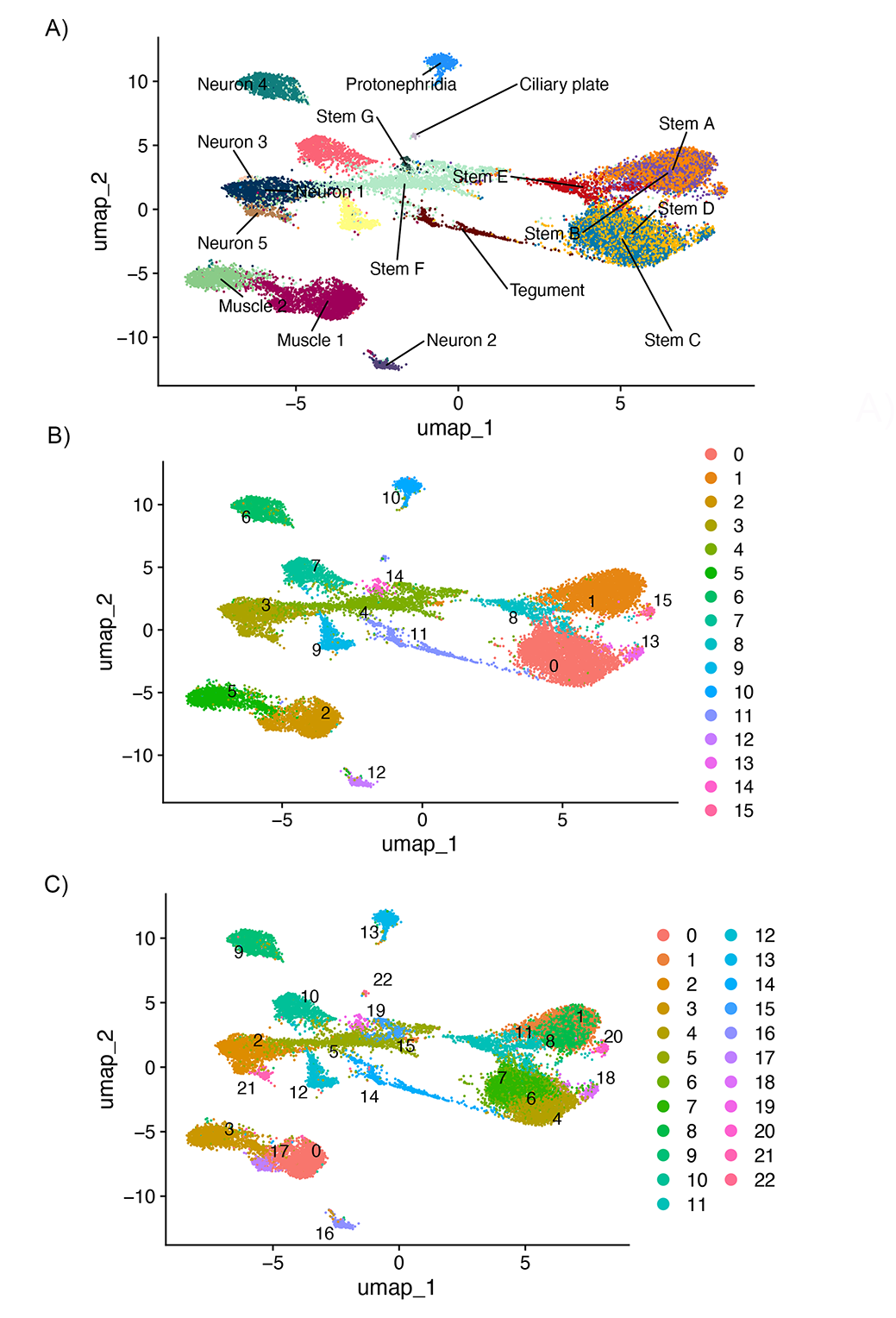

### Supp Fig 23.tif

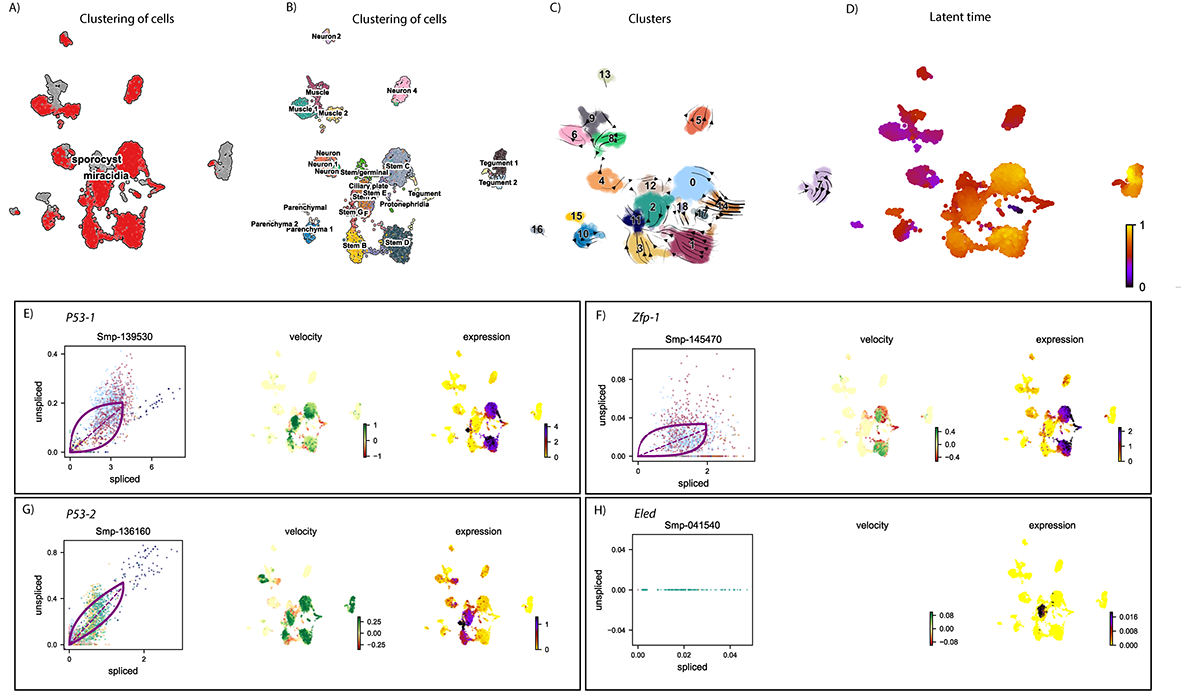
